# Compound activity prediction with dose-dependent transcriptomic profiles and deep learning

**DOI:** 10.1101/2023.08.03.551883

**Authors:** William J. Godinez, Vladimir Trifonov, Bin Fang, Guray Kuzu, Luying Pei, W. Armand Guiguemde, Eric J. Martin, Frederick J. King, Jeremy L. Jenkins, Peter Skewes-Cox

**Affiliations:** Novartis Institutes for BioMedical Research, Emeryville, CA 94608, USA; Novartis Institutes for BioMedical Research, San Diego, CA 92121, USA; Novartis Institutes for BioMedical Research, Cambridge, MA 02139, USA

## Abstract

Predicting compound activity in assays is a long-standing challenge in drug discovery. Computational models based on compound-induced gene-expression signatures from a single profiling assay have shown promise towards predicting compound activity in other, seemingly unrelated, assays. Applications of such models include predicting mechanisms-of-action (MoA) for phenotypic hits, identifying off-target activities, and identifying polypharmacologies. Here, we introduce Transcriptomics-to-Activity Transformer (TAT) models that leverage gene-expression profiles observed over compound treatment at multiple concentrations to predict compound activity in other biochemical or cellular assays. We built TAT models based on gene-expression data from a RASL-Seq assay to predict the activity of 2,692 compounds in 262 dose response assays. We obtained useful models for 51% of the assays as determined through a realistic held-out set. Prospectively, we experimentally validated the activity predictions of a TAT model in a malaria inhibition assay. With a 63% hit rate, TAT successfully identified several sub-micromolar malaria inhibitors. Our results thus demonstrate the potential of transcriptomic responses over compound concentration and the TAT modeling framework as a cost-efficient way to identify the bioactivities of promising compounds across many assays.

## Main

Testing compounds in assays to experimentally ascertain their bioactivity is a core undertaking in drug discovery. Conducting those experiments is a resource-intensive endeavor. Computational models to predict compound activity profiles across many assays have the potential to suggest additional important experiments and to highlight the more promising compounds for follow-up.

Conventional QSAR (quantitative structure-activity models) models mapping compound structure to compound activity have seen limited success, as the models’ predictive performance decreases for compounds with low chemical similarity to compounds in the training set^1,2^. By supplementing the purely structural models with experimental outcomes^3–5^, models have been able to infer activity more accurately for a broader range of assays and chemical matter. Recent developments in transcriptomic profiling technologies^6–10^ have allowed the acquisition of large collections of compound-induced gene-expression signatures that are primed for predictive modeling. Indeed, classification models trained on gene-expression data^11,12^ have been able to predict compound activity in other assays. The combination of transcriptomic and imaging data, together with molecular structure, has also resulted in gains in predictive performance^5,12–15^. Previous models trained on gene-expression profiles have used data from compound treatment at a *single* concentration only. Prior work^16,17^ suggests rather that transcriptomic responses over *multiple* concentrations could lead to improved predictions of compound activity.

This paper introduces transcriptomics-to-activity Transformer (TAT) models that map gene-expression profiles attained over compound treatment at multiple concentrations in a single profiling assay to compound activity in other dose-response assays. We use activity defined as pAC_50,_ the negative logarithm of the half-maximal activity concentration. We trained TAT models on gene-expression data from a RASL-Seq profiling assay^7^ to predict the pAC_50_ values of 2,692 compounds in a panel of 262 dose-response assays. Using a “realistically novel” held-out set^3^, we demonstrate that TAT models provided useful pAC_50_ predictions for 51% of the assays, which substantially surpassed the predictive performance of models trained with single-concentration profiling data. Prospectively, we experimentally validated the predictions of a TAT model in a malaria inhibition assay to show the methodology’s ability to identify sub-micromolar malaria inhibitors.

## Results

### TAT: Transcriptomics-to-activity Transformer

We developed the TAT modeling methodology that takes as input a *multi-concentration transcriptomic profile* as induced by compound treatment at multiple concentrations and measured through a gene-expression profiling assay (**Figure 1**). In such profiling assays, the treatment condition (compound concentration) is replicated multiple times. To build the input multi-concentration profiles for the model, we adopt a combinatorial approach that both augments the data and accounts for the experimental variability observed over the replicates by enumerating all possible paths through a *trellis* entailed by the set of compound concentrations and the set of replicates per concentration (**Figure 2a**). Each path through the trellis represents a multi-concentration transcriptomic profile that includes the gene-expression outcomes at three concentrations for a given compound. Each gene-expression outcome at a certain concentration provides a numerical snapshot of transcriptomic activity through the read counts per gene probe measured at the corresponding concentration replicate (**Methods**). Assuming that all concentration treatments have the same number of replicates, the total number of multi-concentration profiles generated by a single compound through this approach is 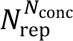, where *N*_conc_ and *N*_rep_ are the number of concentrations and replicates, respectively.

**Figure 1.**
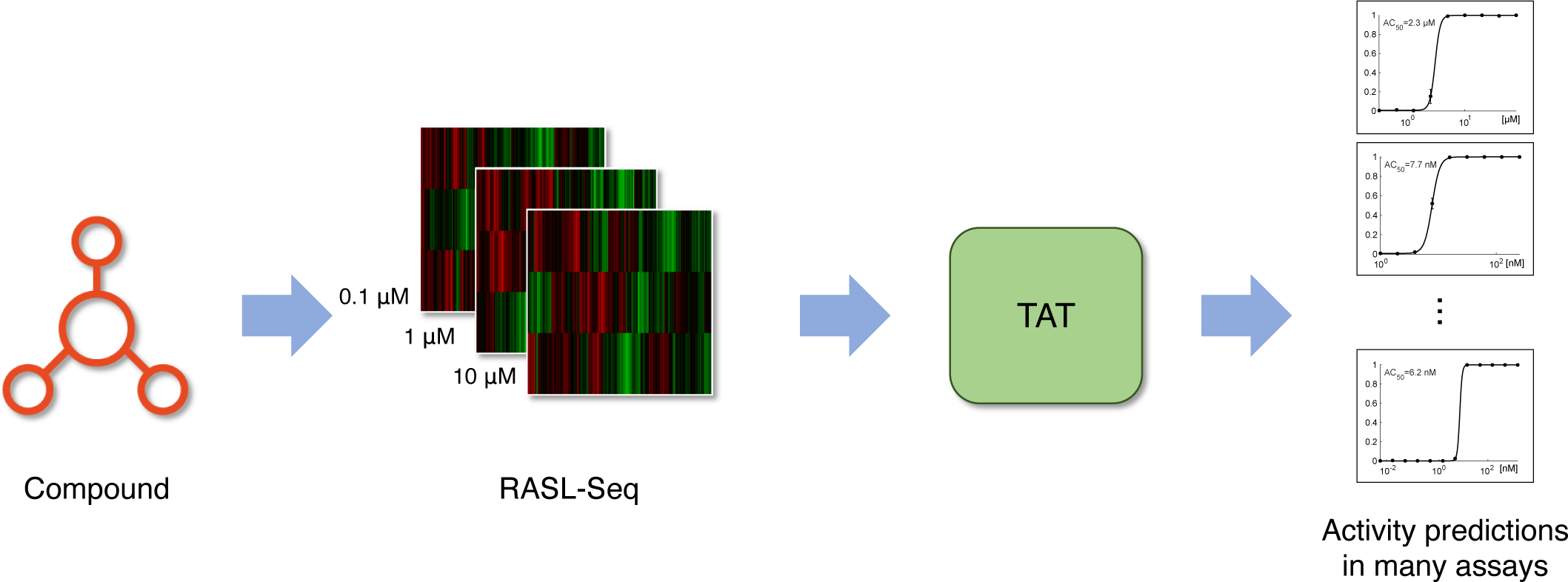
Overview. Transcriptomic outcomes induced by compound treatment at multiple concentrations are measured through a RASL-Seq assay. The resulting multi-concentration transcriptomic profiles serve as input to transcriptomics-to-activity Transformer (TAT) models that translate those profiles into compound activity in other, seemingly unrelated, assays. TAT models are trained one assay at a time.

**Figure 2.**
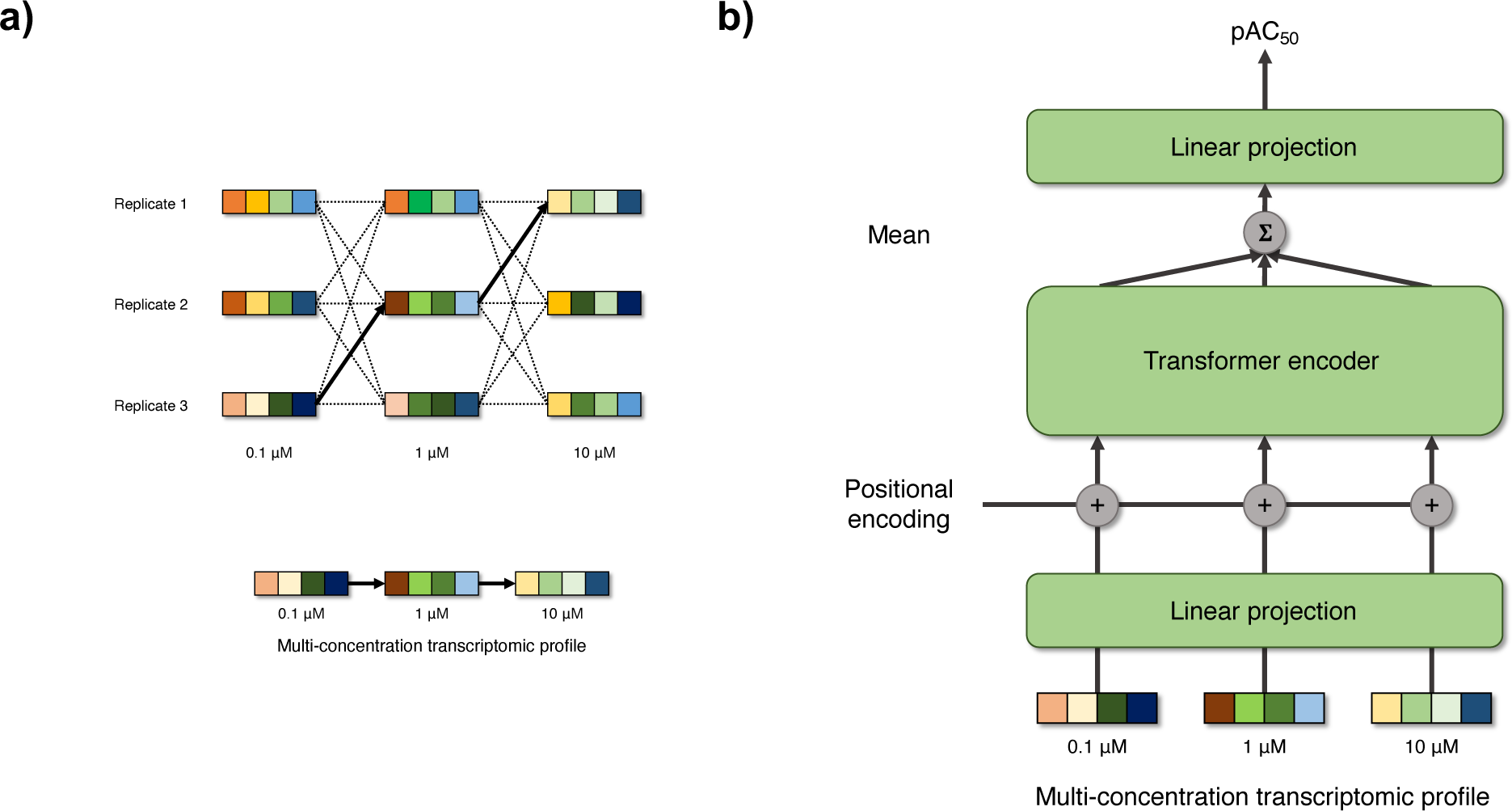
Input and model architecture. **a**) Example trellis diagram representing possible multi-concentration transcriptomic profiles over three concentrations with three replicates each. The multi-colored vectors represent the numerical measurements of transcriptomic activity at a particular concentration replicate. The dashed lines show all 27 possible multi-concentration profiles over the concentrations. The bold arrows in the trellis highlight a particular multi-concentration profile, which is also shown in the bottom as a sequence of transcriptomic outcomes. **b**) Diagram depicting the architecture of the Transcriptomics-to-activity Transformer (TAT) model. Starting from an input multi-concentration profile, the model projects linearly the gene-expression outcomes at individual concentrations, injects positional information encoding the concentration information, and feeds the resulting vectors onto a Transformer encoder. The model then averages the transformed tokens and projects the mean embedding onto a pAC_50_ prediction. One TAT model is trained per assay.

To map the multi-concentration transcriptomic profiles onto pAC_50_ values in an assay of interest, the TAT models rely on the Transformer neural network architecture^18^. Originally designed for natural language processing (NLP) problems dealing with word sequences^19–21^, the Transformer architecture has seen immense success beyond NLP^22–24^, including drug discovery^25–27^, because of its ability to contextualize the long-range correlations among elements of an input sequence through its attention mechanism. Here, the TAT models exploit that mechanism to build multiple hypotheses regarding the importance of gene-expression outcomes at individual compound concentrations relative to the compound’s activity in an assay of interest. Architecturally, the model first projects linearly the gene-expression outcomes at individual concentrations in an input multi-concentration profile onto a learned 512-D feature space. After injecting the positional information that implicitly encodes the compound concentration, the resulting concentration-informed gene-expression embeddings are fed to a single encoder block of the Transformer. The transformed embeddings (tokens) are then averaged to obtain a single 512-D vectorial representation that subsumes the information in the input multi-concentration transcriptomic profile. The 512-D encoding is projected linearly to predict a pAC_50_ value of a compound in an assay of interest (**Figure 2b**). TAT models are trained by optimizing a mean squared error (MSE) criterion that compares a model’s pAC_50_ predictions with experimental pAC_50_ measurements (**Methods**).

### TAT models predict successfully a majority of panel of assays

To validate the TAT methodology, we used 2,692 compounds of the Novartis Mechanism-of-Action (MoA) Box chemogenetic library^28^ that were profiled in a RASL-Seq gene-expression assay. In this assay, U-2 OS cells were compound-treated in triplicate at three concentrations. A collection of genes based on the L1000 gene set^8^ was used to monitor transcriptomic activity in the cell line (**Methods**). Using our combinatorial approach to assemble the input data for the TAT model, we obtained 27 multi-concentration profiles per compound, for a total of 72,684 profiles.

Subsets of these 2,692 compounds had been tested and shown activity in 262 biochemical and cellular dose-response assays. This diverse set of assays includes a variety of cell lines as well as biochemical assays from diverse target classes. The size of the subsets tested in those assays ranged from 403 compounds up to 2,248 compounds (**Supplementary Table 1**).

We proceeded to examine the ability of TAT models to predict compound activity in each of the 262 assays. For benchmarking purposes, we also evaluated the performance of a random forest (RF) model as well as a standard feed-forward neural network (NN) model on each assay. For the latter two models, we used the concatenation of gene-expression outcomes at individual concentrations within each multi-concentration profile as input feature vectors. As an example, the concatenation of the three vectorial elements of the multi-concentration profile in **Figure 2a** constitutes the input feature vector to the RF and NN models. We also built RF models using gene-expression profiles derived from treatment at single compound concentrations, in which case we used the element-wise mean profile over the concentration replicates as input, which is standard practice in the literature^11–13,29^. In total, we evaluated six models per assay. All parameters for all models were held constant across assays (**Methods**).

To evaluate the performance of the models, we separated the compounds in each assay into a training set and test set through a “realistically novel” split that assigns into the test set chemical structures that are uncommon (i.e., ‘novel’) relative to the compound collection profiled in the assay^3^. To numerically determine the performance of a model in a single assay, we calculated the squared Pearson correlation coefficient *r*^2^ between predicted and experimental pAC_50_ values on the realistic test set. To ascertain the performance across the panel of assays, we counted the number of assays for which each model exceeded a value of *r*^2^ = 0.3 on the test set as defined in prior work^3^.

The cumulative distributions of *r*^2^ values for each model are shown in **Figure 3**. The upper three curves correspond to RF models built with single concentration data, denoting that these models predominantly yielded lower *r*^2^ values. Median *r*^2^ values for these models ranged from 0.1 to 0.17 and 14–23% of the models exceeded our threshold of success *r*^2^ > 0.3 (**Table 1**). As hypothesized, the accuracy of the RF model is substantially improved by using gene-expression profiles from all concentrations, with the model achieving a median *r*^2^ value of 0.23 and successfully modeling 39% of the assay collection. A nominally better performance is obtained by the NN model, with a median *r*^2^ value of 0.24 and 41% assays modeled successfully. Unlike the RF and NN models, the TAT model additionally leverages the context afforded by the gene-expression outcomes encoded in the multi-concentration transcriptomic profile. It thus achieved an improved median *r*^2^ of 0.31 and is the only approach to successfully model a majority of assays (51%).

**Figure 3.**
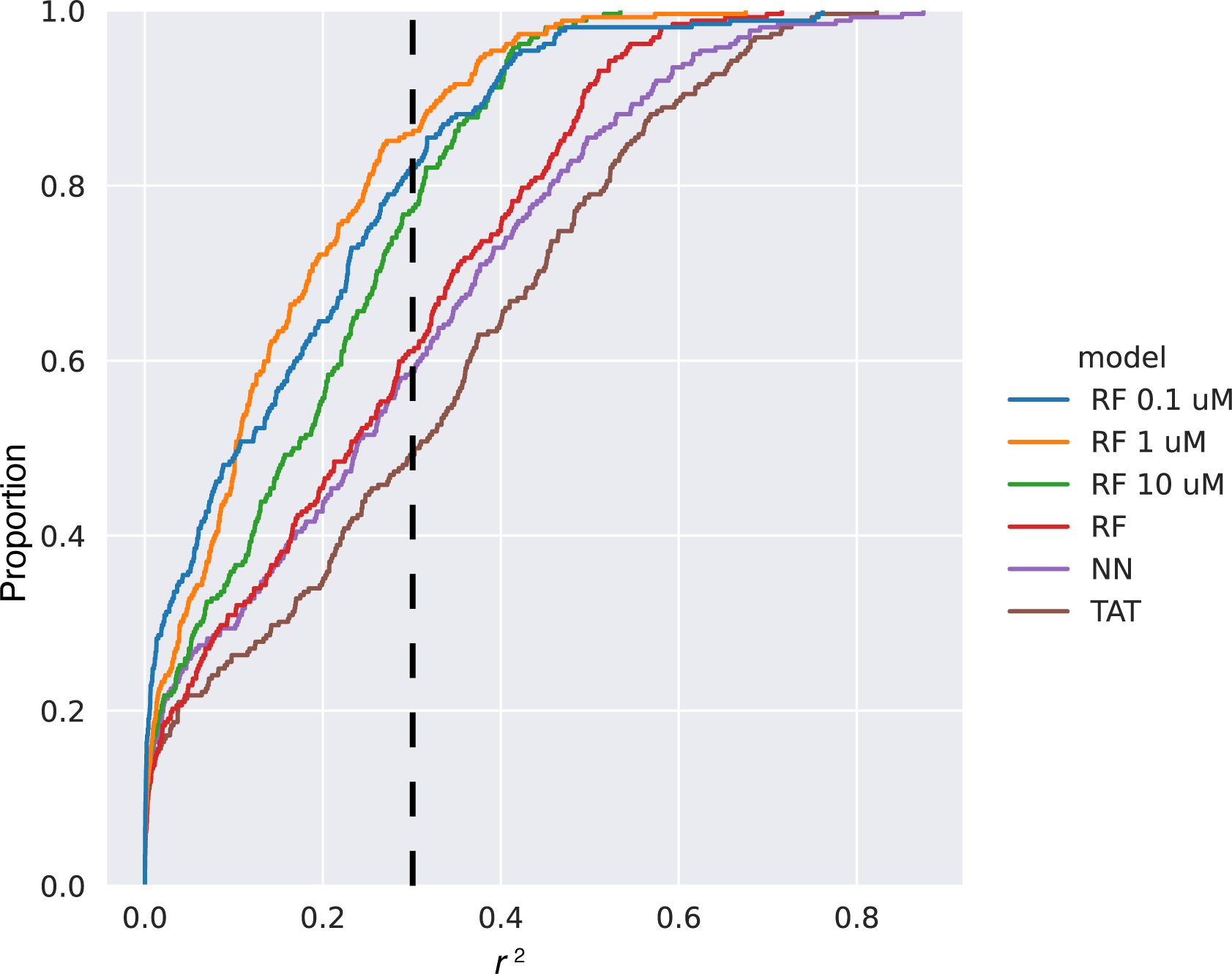
Cumulative distribution of *r*^2^ values for models across a panel of 262 assays. We show the performance of models built using single concentration data (RF 0.01-10 µM) as well as models built with all concentration data (NN, RF, and TAT). The vertical black dashed line highlights the *r*^2^ > 0.3 success threshold. RF – Random Forest, NN – Neural Network, TAT – Transcriptomics-to-Activity Transformer.

**Table 1.** Performance of models built with RASL-Seq data across a panel of 262 assays. The table shows the number and percentage of assays successfully described by each approach (model-data combination).

| **Model** | **Concentration data** | **Median $r^2$** | **Number of successfully modeled assays** | **Percentage of successfully modeled assays ($r^2>0.3$) [%]** |
| --- | --- | --- | --- | --- |
| **RF** | 0.1 μM | 0.11 | 47 | 18 |
| **RF** | 1 μM | 0.10 | 37 | 14 |
| **RF** | 10 μM | 0.17 | 60 | 23 |
| **RF** | All | 0.23 | 102 | 39 |
| **NN** | All | 0.24 | 108 | 41 |
| **TAT** | All | 0.31 | 133 | 51 |

TAT successfully modeled various cellular assays, including cell proliferation assays, cell viability assays, and toxicity assays (**Figure 4a**). While TAT is trained on gene-expression readouts from a U-2 OS cell line, it still managed to predict activity outcomes in other cell lines (**Supplementary Figure 1**). Biochemical assays monitoring compound activity against certain target classes, such as kinases and GPCRs, were also predictable from gene-expression profiles through TAT (**Figure 4b**). The distribution of activity of compounds in the assays, as reflected by the standard deviation of pAC_50_ values in each assay, bore no correlation (r^2^ < 0.001) with the ability of TAT to successfully model an assay (**Supplementary Figure 2**). Model performance rather appears to be related to the biology captured by individual assays. For example, receptor internalization assays could not be predicted, even when these assays involved the same cell line (U-2 OS) used to train TAT, presumably because receptor internalization activity is measured over a short time scale (e.g., minutes), while the measurements of transcriptomic activity used to train TAT were captured 24 hours after treatment.

**Figure 4.**
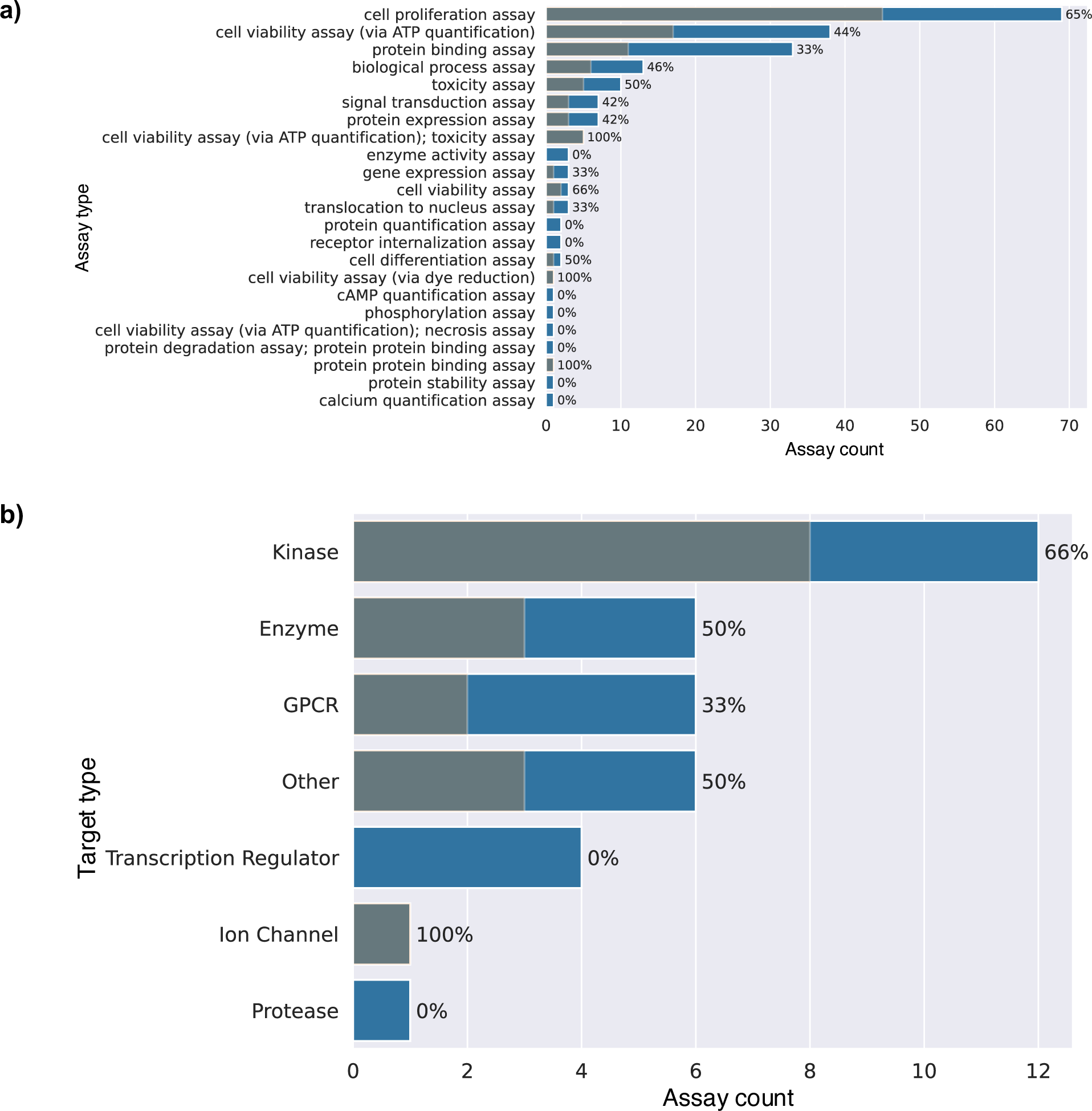
Number of assays successfully modeled by TAT. The blue bars show the number of assays in each assay category, while the stacked gray bars show the number of assays successfully modeled by TAT within each category. The percentage of successful models is listed next to each bar. The entries are sorted by the number of assays in each category. **a**) Number of successfully modeled assays per assay type. **b**) Number of successfully modeled assays per target type for target-based assays.

### TAT prospectively identifies compounds with sub-micromolar malaria inhibition activity

Having computationally ascertained the ability of TAT models to provide useful predictions on existing assay data, we proceeded to experimentally validate the model’s prospective predictions in a Novartis-internal *Plasmodium falciparum* inhibition assay (see **Methods**). This assay was not included in the previous collection of 262 assays. 482 compounds had been both tested in the malaria inhibition assay and profiled in RASL-Seq. The compounds were clustered and a TAT model was trained using 75% of those compounds from the largest clusters. The correlation of prediction with experiment for the remaining 25% “realistic” test set of singletons and small clusters was *r*^2^ = 0.33, meeting our standard for success of *r*^2^ > 0.3, as well as demonstrating the ability of TAT to predict compound activity in a non-human cell line.

We rebuilt the TAT model with all 482 compounds (no test set withheld) and predicted pEC_50_ values for the 3,256 compounds for which we had RASL-seq data but had not been tested in the malaria inhibition assay. To determine whether the model would be useful for identifying active compounds, we picked the top predicted 1% of compounds for experimental testing (*n* = 33, three compounds were unavailable). To estimate the background hit rate of this list of 3,256 compounds, we made a random selection of 33 compounds. We also picked the bottom 1% of the list of compounds (*n* = 33) to see whether the model predictions for the compounds predicted to be most inactive would be accurate. All 96 compounds were then tested in vitro against the 3D7 strain of *P. falciparum*. For these compounds, the correlation of prediction with experimental pEC_50_ was *r*^2^ = 0.52, the Spearman rank correlation was ρ^(^ = 0.53, and the mean absolute error (MAE) was 0.39 (**Figure 5a**). Using a typical hit threshold of EC_50_ < 1 μM (pEC_50_ > 6), there were no hits in the 1% predicted least active (**Table 2**). Likewise, the random category had no hits, indicating that the background hit-rate is < 3%. For the active category, 19 out of 30 compounds exhibited activity below 1 μM—a 63% hit rate. So the false-negative rate was very low for our sampling size, and the false positive rate was only 37%.

**Figure 5.**
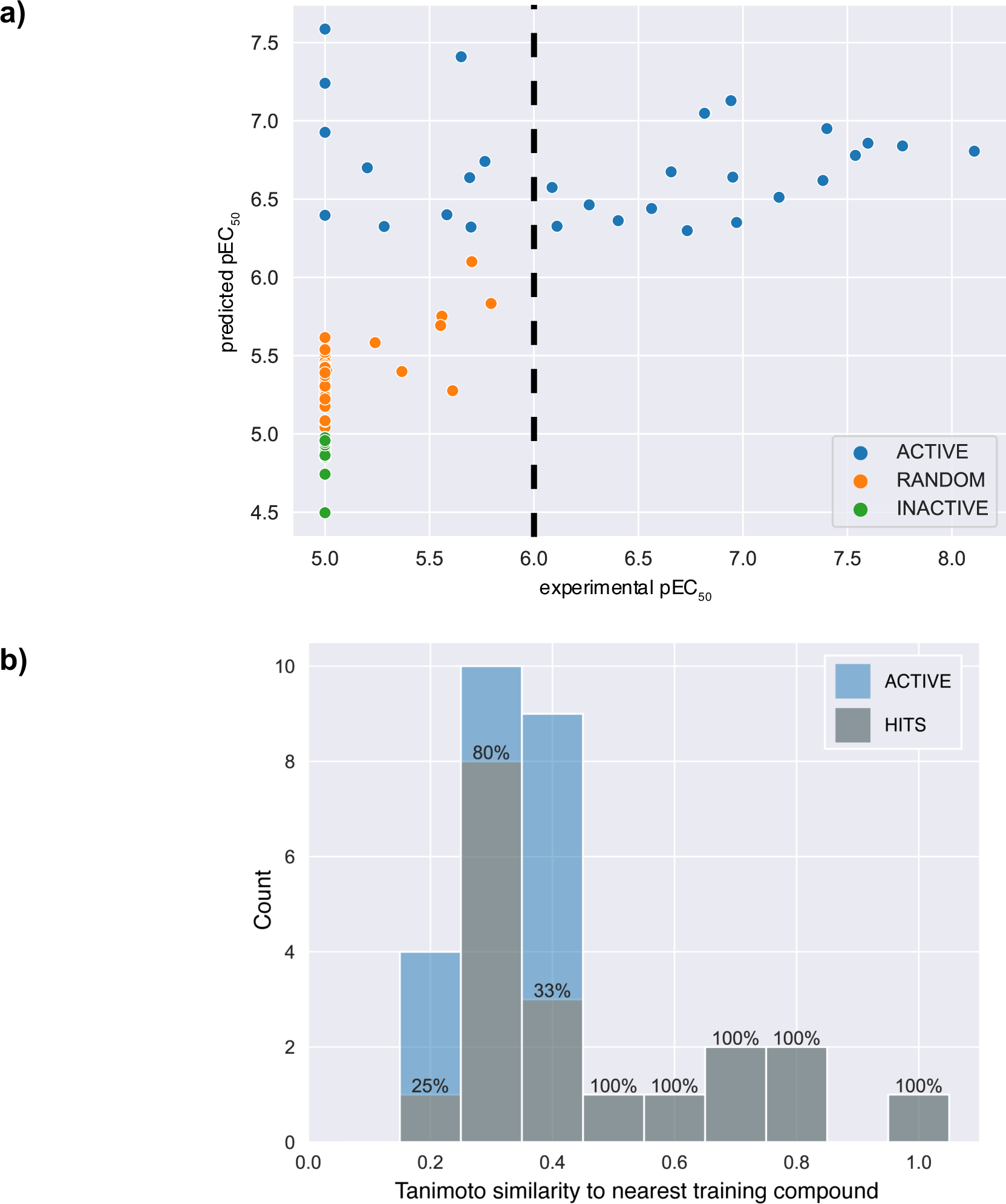
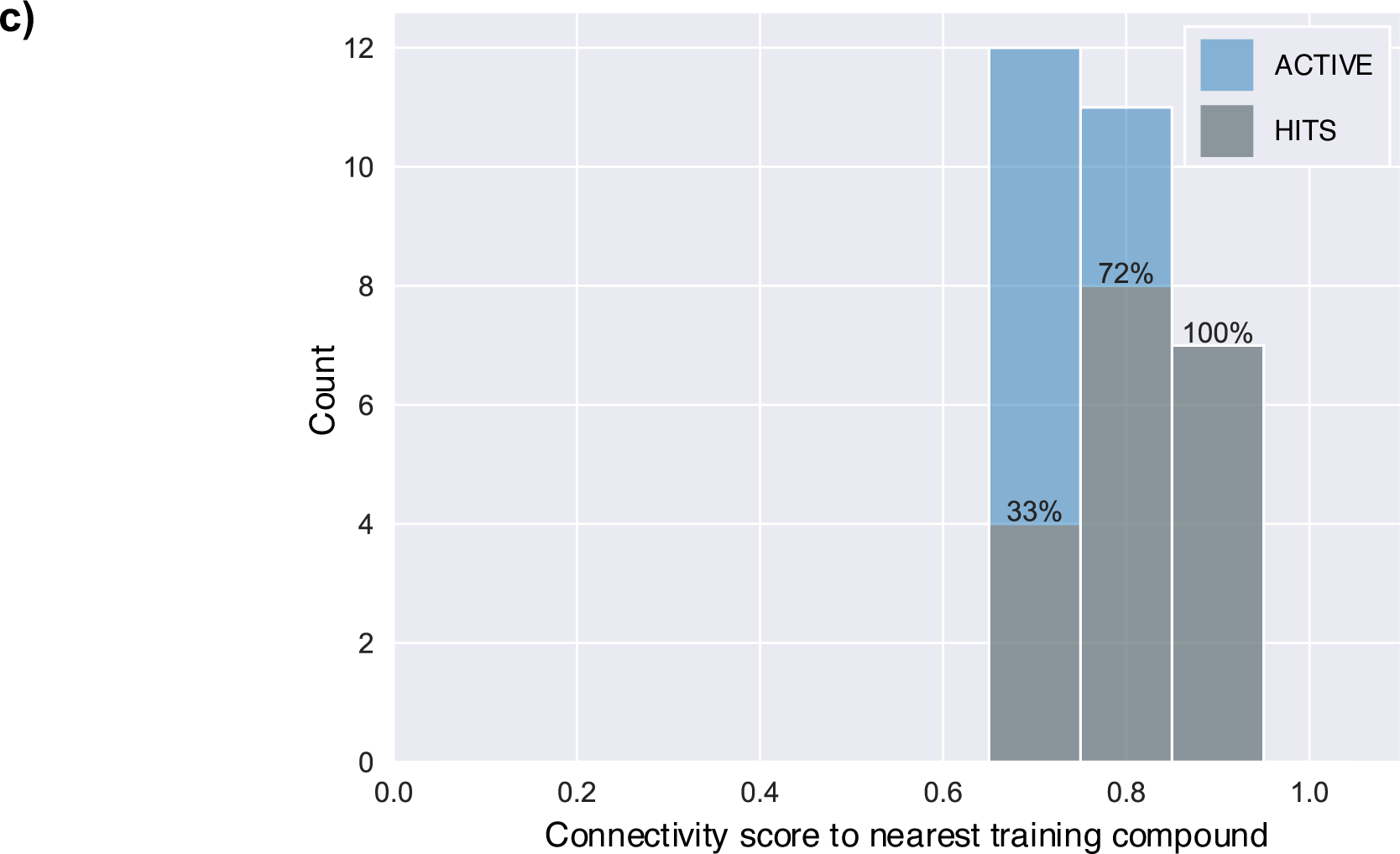
Experimental results for compounds picked by a TAT model in a malaria inhibition assay. **a)** Experimental pEC_50_ values (x-axis) versus the model’s pEC_50_ values (y-axis). Each data point represents a compound. The color for each data point represents the category (active, random, or inactive) for the corresponding compound, as determined through the model pEC_50_ prediction. The vertical black dashed line highlights the EC_50_ < 1 μM hit threshold (pEC_50_ > 6) at which a compound is considered to be active. **b**) Blue bars depict the histogram of Tanimoto coefficients for compounds (n=30) predicted to be active by the model to their nearest neighbor compound in the training set. Stacked grey bars show the distribution of Tanimoto coefficients for the subset of compounds (n=19) that were both predicted to be active by the model as well as active in the assay (true positives or hits). Hit rates are shown on top of each Tanimoto coefficient bin. **c**) Histogram of connectivity scores for compounds (n=30) predicted to be active by the model to their nearest neighbor compound in the training set. Stacked grey bars show the distribution of connectivity scores for the subset of compounds (n=19) that were both predicted to be active by the model as well as active in the assay (true positives or hits). Hit rates are shown on top of each connectivity score bin.

**Table 2.** Number of compounds, hits and hit rate for the different pick categories in a malaria inhibition assay.

|  | INACTIVE | RANDOM | ACTIVE |
| --- | --- | --- | --- |
| **Compounds** | 33 | 33 | 30 |
| **Hits (< 1 μM)** | 0 | 0 | 19 |
| **Hit Rate** | 0% | 0% | 63.33% |

We also investigated the ability of the TAT model to pick novel active chemical matter. For each compound in the active pick category, we computed the Tanimoto coefficient (Tc) for similarity between each compound predicted as active and its nearest neighbor in the training set. The stacked histogram of Tc of the predicted actives and hits to their corresponding nearest neighbor in the training set is shown in **Figure 5b**. Most of the predicted actives had very low chemical similarity to the training set. The hit rates calculated for each similarity bin show 100% hit-rates for Tc > 0.4, and hit-rates from 25– 80% even for novel compounds with Tc < 0.4. We also calculated the hit rates as a function of the connectivity scores between the profiles of compounds predicted to be active and their nearest neighbor molecule, in terms of connectivity score, in the training set (**Figure 5c**). TAT managed to pick hits even for compounds with relatively different transcriptomic profiles (connectivity score < 0.8).

## Discussion

The development of high-throughput gene-expression profiling technologies^6–10^ has enabled the acquisition of large collections of compound-induced transcriptomic signatures. These large collections provide novel modeling opportunities for target identification^30,31^ and compound bioactivity prediction^11–13^. Current transcriptomic-based approaches for compound activity prediction, which rely on gene-expression readouts induced by compound treatment at a single concentration, model only a limited number of assays. Our TAT machine learning models that leverage transcriptomic readouts induced by compound treatment at multiple concentrations successfully model substantially more assays.

TAT tackles the activity prediction task using a Transformer neural network model that contextualizes the relative importance of gene-expression profiles at different concentrations. TAT models are trained with a novel data augmentation scheme that exploits the granularity and variability afforded by the experimental replicates in the gene-expression profiling experiment. This stands in contrast with methodologies that aggregate over replicates^11–13^.

We validated computationally the performance of TAT models using a collection of 262 assays that included a diversity of assay modalities, cell lines, and targets. TAT was especially successful for cellular assays, including cellular proliferation, cell viability, and toxicity assays. While TAT was trained on transcriptomic activity from an osteosarcoma cell line (U-2 OS), it managed to predict activity in other cell lines. Overall, TAT successfully predicted a majority of the assays in the collection.

Prospectively, we used the TAT model to identify novel active compounds in a malaria inhibition assay. With a hit rate of 63% (over an estimated background hit rate of <3%) and no false negatives, we showed the potential gains in experimental efficiency that a TAT model could bring to a drug discovery project. Through an evaluation of the chemical similarity between the model’s active picks and compounds in the training set, our prospective study also highlighted the ability of TAT to identify novel potent chemical matter.

TAT is currently trained on transcriptomic data measured through RASL-Seq. This targeted sequencing technology allows us to obtain, in a cost-effective manner, robust numerical snapshots of cellular activity that can be readily fed to machine learning models. Conceptually, TAT is suitable for modeling other sequencing technologies, such as DRUG-Seq^10^, as well.

Given its potential, we envision a few use cases where RASL-Seq together with TAT could impact drug discovery. As demonstrated through the application to the malaria assay, RASL-Seq and TAT can serve as an accurate proxy for an individual assay. Expensive or low throughout assays would especially benefit from this aspect of the methodology, as RASL-Seq and TAT modeling could provide an alternative for anticipating compound activity in such challenging assays. In particular, for in vivo assays, the TAT models could flag up compound liabilities early in the drug discovery pipeline and thus help prioritize lead compounds for more expensive in vivo testing. The ability of TAT to predict activity in historical assays also makes it helpful in potentially repurposing drugs for new indications. The proposed approach also provides improvements in experimental efficiency through its wholesale predictions across scores of assays, as it is more practical to run a single RASL-Seq assay and TAT modeling instead of running hundreds of individual assays. The predicted activity profile of compounds across scores of assays can provide insights on the compounds’ potential targets, off-targets, mechanism-of-action, and poly-pharmacology.

As TAT is trained on transcriptomic activity from a single cell line as monitored through a fixed gene panel and recorded at a single time point, it is plausible that certain cell-specific mechanisms manifesting themselves at varying time points might not be captured through our current transcriptomic measurements. The inclusion of additional cell lines, time points, and gene probe sets might improve^32^ the modeling scope of TAT and remains the subject of future work.

In conclusion, our study shows the broad potential of transcriptomic data and machine learning for distinguishing the more promising chemical matter early on in the drug discovery pipeline. Application of the TAT methodology to a variety of assays is relatively straightforward. Conceptually, TAT could also be trained on imaging profiles of phenotypes induced by compounds in a series of concentrations. Future work involves algorithmic developments to impute computationally gene-expression data from compound structure to expand the scope of chemical applicability of the models.

## Acknowledgments

The authors thank all people at Novartis who gathered the experimental assay data used in this study. We also thank Chris Roberts, Frantisek Supek, and Loren Miraglia for insightful discussions.

## Author contributions

W.J.G. designed and led the study. W.J.G. developed, implemented, and evaluated TAT. V.T., B.F., and G.K. performed bioinformatics modeling and provided feedback. L.P. conducted the malaria experiments and collected data. F.J.K. led the gene expression profiling experiments. P.S.-C., W.A.G., F.J.K., and J.L.J., provided computational and experimental resources as well as feedback. W.J.G., E.J.M., B.F., W.A.G., and P.S.-C. analyzed and interpreted the results. W.J.G. and E.J.M. wrote the article. All authors reviewed the manuscript.

## Competing interests

All authors are (or were at the time of their involvement with the studies) employees of Novartis.

## Additional information

Supplementary information for this paper is available in **Supplementary Figures** and **Supplementary Tables.**

## Data availability

The data used in this study is proprietary to Novartis. The data is not publicly available due to intellectual property restrictions. An example dataset is available from the Broad Institute at https://broad.io/rosetta/.

## Code availability

The code for TAT is available in **Supplementary Code** and at https://github.com/Novartis/TAT

## Methods

### RASL-Seq assay

Using a published protocol^7^, the RASL-Seq assay quantitates the expression of 978 genes in U2-OS (American Type Culture Collection (ATCC), cat. no. HTB-96) samples. For each gene, three RASL-seq probe sets were designed and incorporated into the analysis. Samples were exposed for 24 hours to small-molecule treatment before measuring transcriptomic activity. Each treatment was performed in triplicate.

### Dataset

In this study, we used 2,692 compounds from the Novartis Mechanism-of-Action (MoA) Box chemogenetic library^28^. These compounds were profiled in triplicate through the RASL-Seq gene expression assay at three concentrations: 0.1 µM, 1 µM, and 10 µM. For each replicate, the RASL-Seq assay yields numerical readouts for three probes per gene. By concatenating the three gene probes, we obtained a 2934-D gene-expression profile per replicate. The gene counts for each replicate are transformed to reads-per-million (RPM) and normalized on a plate-by-plate basis using the RPM values of the plates’ neutral control treatments (DMSO).

### Multi-concentration transcriptomic profiles

The Transcriptomics-to-activity Transformer (TAT) takes as input a multi-concentration gene expression profile. To assemble those multi-concentration profiles per compound, we define a trellis as a *k*-partite graph, with *k* defining the number of independent sets in the graph. In our case, *k* is given by the number of available compound concentrations *N*_conc_ at which gene-expression profiles have been measured. The vertices in the *c*-th set of the graph are given by the gene expression profiles 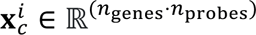 where *c* ∈ *C* is the concentration index, *C* is the linearly ordered set of available concentration indices, *i* ∈ *P* is the replicate index, *P* represents the set of replicate indices, *n*_genes_ is the number of genes monitored, and *n*_probes_ is the number of probes measured per gene. We assume that the number of replicates *N*_rep_ is constant over all concentrations, and therefore the number of vertices in each independent set in the graph is |*P*| = *N*_rep_. Edges in the graph go from all vertices in set *c* to all vertices in set (*c* + 1). An example trellis for *N*_conc_ = 3 and *N*_rep_ = 3 is shown in **Figure 2a**. By enumerating all possible paths starting at a vertex at the lowest concentration set and terminating at a vertex at the highest concentration set, we obtain sequences 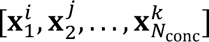 of replicate profiles over concentration, where *i*, *j*, …, *k* ∈ *P*. Per compound, we obtain a total of 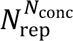 of such paths, with each path representing a multi-concentration transcriptomic profile for that compound. The dataset used in this study included *N*_conc_ = 3 concentrations per compound and *N*_rep_ = 3 replicates per concentration leading to 27 multi-concentration transcriptomic profiles per compound.

### Model Architecture

An overview of the model is shown in **Figure 2b**. The Transcriptomics-to-Activity Transformer (TAT) model aims to learn a mapping between a multi-concentration transcriptomic profile of a compound and that compound’s pAC_50_ in an assay of interest. The TAT model uses only the encoder section of the Transformer architecture^18^. The input to the Transformer encoder is a sequence of gene-expression profiles 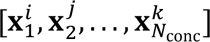 as defined in the previous section. The number of concentrations *N*_conc_ becomes the input sequence length for the Transformer encoder. The gene-expression profiles 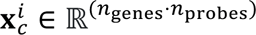 are each encoded onto a feature space ℝ^D^ with a learned linear projection 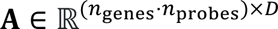. Positional embeddings ***p****_c_* ∈ ℝ^D^, as detailed in the original Transformer article^18^, are added to the projected gene expression profiles 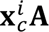 to introduce the relative position of each profile as entailed by the corresponding concentration. The resulting concentration-aware gene-expression encodings 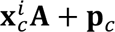 are passed altogether onto a single Transformer encoder block *TE*(·)^18^ that residually integrates information calculated by a multi-head self-attention module as well and a multi-layer perceptron module. The input and output vectors in the encoder modules are of dimension *D*. The resulting *D*-dimensional tokens 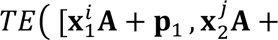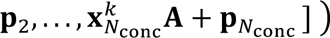 are averaged onto a single vector ***z*** ∈ ℝ^D^. The latent representation **z** subsuming all information in the multi-concentration profile is finally linearly projected onto the output pAC_50_ value. In the TAT models built for each individual assay, we set *D* = 512 as in the original Transformer article^18^, and use four self-attention heads for the Transformer encoder.

### Training

To train the model for an assay of interest, we use the multi-concentration transcriptomic profiles of a set of compounds along with the compounds’ experimental pAC_50_s in the assay of interest. During training, the goal is to improve the ability of the model to predict a pAC_50_ value given a multi-concentration gene expression profile by minimizing the following loss function:

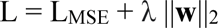

where L_MSE_ is the mean square error (MSE) between predicted and experimental pAC_50_s, ||**w**||_2_ is the L2 norm of the vector **w** including all model parameters, and λ is the weight decay coefficient. For optimization, we use the “noam” scheme as detailed in the original Transformer article^18^. We use a starting learning rate of 0.0 with a warmup rate of 3 epochs and a scale factor of 0.05. For the underlying ADAM optimizer^33^, we use β_1_ = 0.9, β_2_ = 0.98, a batch size of 180, and a weight decay coefficient of λ=0.0005. The dropout rate is set to 0.0. We additionally apply gradient clipping at a global norm of 1. A TAT model on a single assay is trained for 36 epochs. We use PyTorch to implement the model and used an NVIDIA Tesla V100 GPU with 16GB of memory to perform the calculations.

### Model validation

To evaluate the TAT model in a single assay of interest, we split the compounds tested in the assay of interest into a training set and a “realistically novel” hold-out set^3^. The realistic approach first clusters the compounds based on their chemical similarity. Compounds in the larger clusters are progressively added to the training set until 75% of the compounds are included. The remaining 25% from the smaller clusters and singletons are assigned to the test set. The model is trained on all multi-concentration profiles of all compounds in the training set. Each multi-concentration transcriptomic profile in the training set, together with the pAC_50_ associated with the respective compound in the assay of interest, is treated as an independent training example. At test time, the models produce one pAC_50_ per input multi-concentration transcriptomic profile, thus affording an ensemble of pAC_50_ predictions per compound from which we derive a mean pAC_50_ value that serves as the actual model prediction. The predicted mean pAC_50_ of the compounds in the test set are then compared with the ground truth pAC_50_ values to compute the squared Pearson correlation coefficient *r*^2^ that summarizes the performance of the model in an individual assay. Models achieving a correlation of *r*^2^ > 0.3 are deemed as successful^3^.

### Baseline models

In each assay, the predictive performance of the TAT model is compared to the performance of a random forest (RF) and a feed-forward neural network (NN). For both models, the input is a concatenation 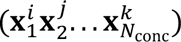 of the gene-expression profiles included in a multi-concentration transcriptomic profile. The number of training examples is therefore the same as the number of training examples used to train a TAT model. Both models are likewise optimized by minimizing a mean-squared error (MSE) criterion. For the RF model we use the Scikit-learn^34^ implementation with 300 trees. The NN model includes four modules, each module consisting of a linear projection layer, a batch normalization layer, a leaky ReLU layer, and a dropout layer. A final output layer projects the last hidden representation to a pAC_50_ value. All hidden layers in the network use 2048 hidden units. The dropout rate is set to 0.1. The network is trained with an ADAM optimizer^33^ for 36 epochs with a batch size of 180 training examples. The starting learning rate is set to 0.0003 and is annealed with an exponential parameter of *γ* = 0.9.

For each assay, we also built random forest (RF) models using profiles from individual concentrations. The input to each model is the mean of the gene expression profiles over replicates at concentration index 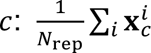. In total we evaluate three models using the data from three concentrations, respectively: 0.1 µM, 1 µM, and 10 µM. The single-concentration RF models are likewise set to use 300 trees.

### 3D7 malaria inhibition assay details

Following an established protocol^35^, *P. falciparum* 3D7 (BEI, MRA-102) parasite cultures were grown with complete media (RPMI 1640 medium (10.4 g/l) with 0.5% AlbuMAX II, 200 µM hypoxanthine, 50 mg/L gentamicin sulphate, 35 mM HEPES, 2.0 g/L sodium bicarbonate and 11 mM glucose) and human erythrocytes. Cultures were maintained at 37° C in an incubator with 5% CO_2_. In vitro antimalarial activity was measured according to a modified SYBR Green cell proliferation assay^36^. Dose response curves data were normalized based on fluorescence signal values from DMSO treated wells (0% inhibition) and mefloquine treated wells (100% inhibition) at a final concentration of 10 µM. The standard four parameter logistic regression model was applied for curve fitting in order to determine EC_50_ using an in-house data analysis system Helios^37^, based on one independent experiment each performed in duplicate.

## Supplementary Figures

**Supplementary Figure 1.**
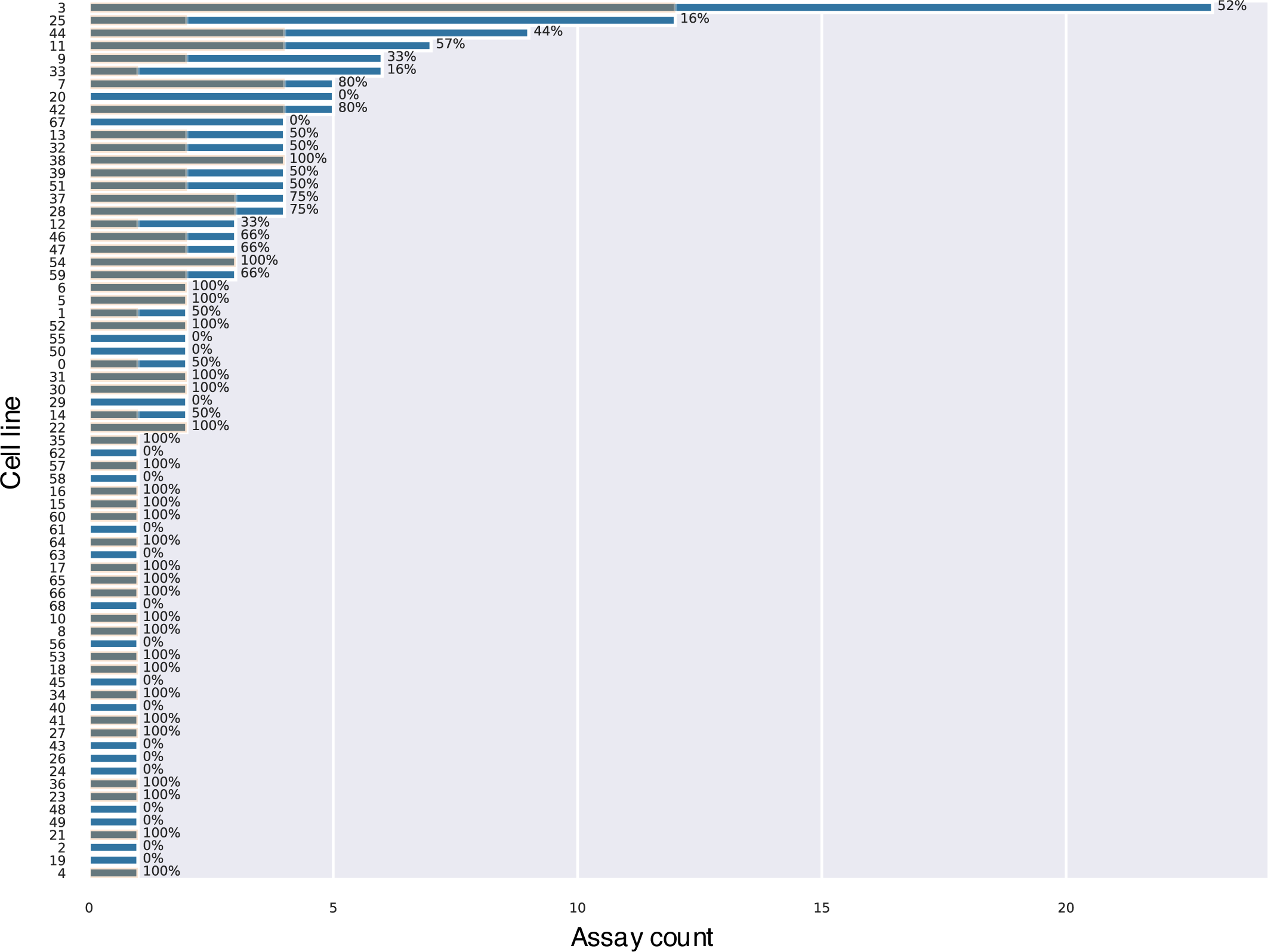
Number of assays successfully modeled by TAT per cell line. The blue bars show the number of assays in each cell line, while the gray overlaid bar shows the number of assays successfully modeled by TAT in a particular cell line. The percentage of successful models is listed next to each bar. The entries are sorted by the number of assays in each cell line. The U-2 OS cell line is labeled as cell line 33.

**Supplementary Figure 2.**
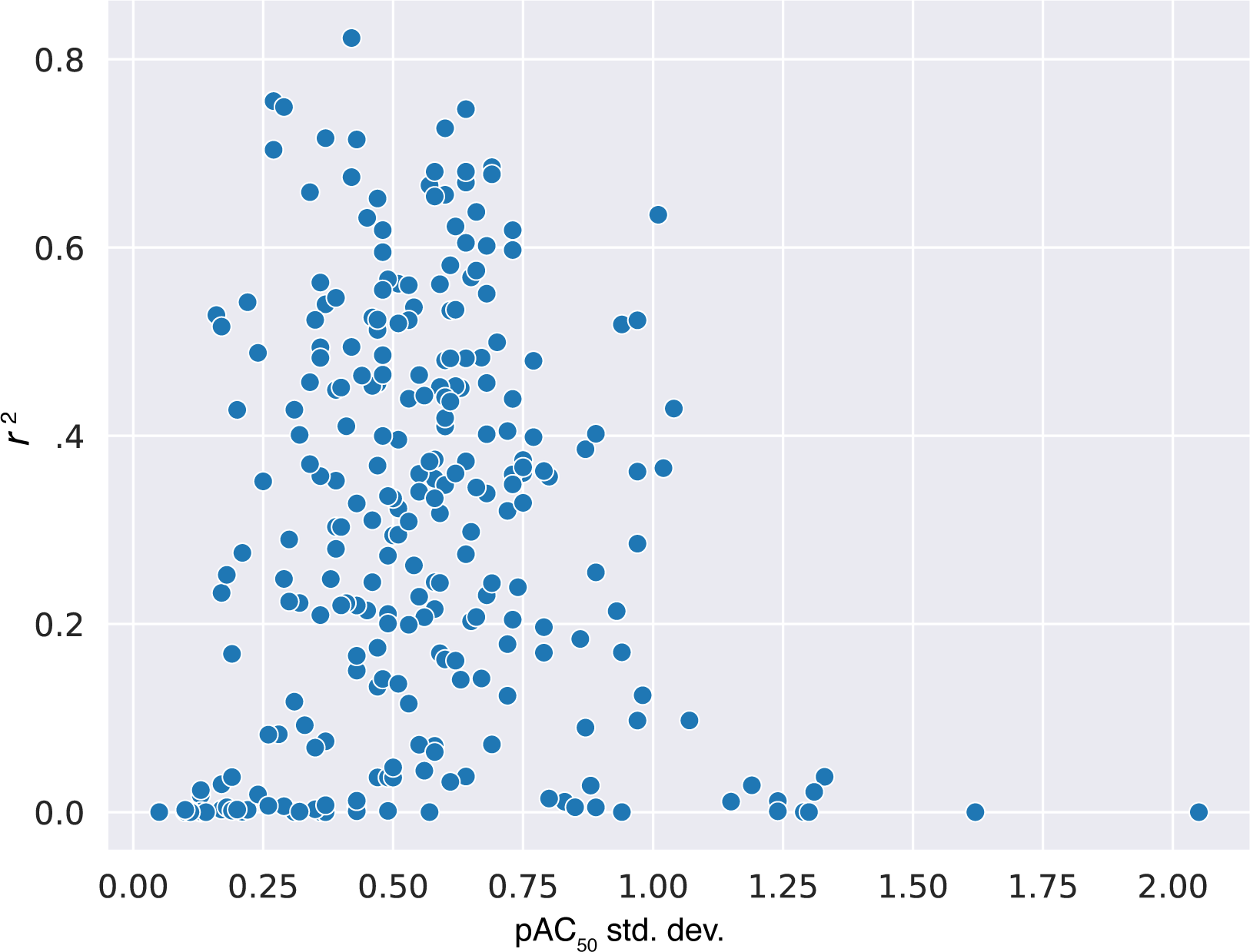
Scatterplot displaying the relation between the standard deviation of pAC_50_ values in each assay (x-axis) and the accuracy of the TAT model, in terms of r^2^ (y-axis).

