## Supplementary Table 1 for "Compound activity prediction with dose-dependent transcriptomic profiles and deep learning"

**Supplementary Table 1**
Description of the 262 dose-response assays involved in our study. Values in the cell line column represent different cell lines, for a total of 69 different cell lines. Cell line index 33 corresponds to the U-2 OS cell line. Other cell lines are not disclosed because of intellectual property restrictions.

| Assay ID | No. of compounds | No. of active compounds | Cell line | Assay type | Target class |
|---|---|---|---|---|---|
| **0** | 2248 | 36 | -- | biological process assay | Protease |
| **1** | 2248 | 149 | -- | | -- |
| **2** | 2248 | 153 | -- | | -- |
| **3** | 2248 | 191 | -- | | -- |
| **4** | 2248 | 1420 | -- | cell viability assay (via ATP quantification) | Kinase |
| **5** | 2248 | 253 | -- | | -- |
| **6** | 2248 | 89 | -- | biological process assay | Other |
| **7** | 2248 | 38 | -- | biological process assay | -- |
| **8** | 2247 | 120 | 68 | biological process assay | -- |
| **9** | 2245 | 11 | 2 | cell viability assay (via ATP quantification); necrosis assay | -- |
| **10** | 2017 | 7 | 25 | cAMP quantification assay | -- |
| **11** | 2017 | 18 | 62 | enzyme activity assay | Enzyme |
| **12** | 1928 | 133 | 59 | | -- |
| **13** | 1928 | 110 | 59 | cell viability assay | -- |
| **14** | 1900 | 65 | -- | | -- |
| **15** | 1900 | 187 | 63 | protein quantification assay | -- |
| **16** | 1900 | 8 | -- | protein degradation assay; protein protein binding assay | -- |
| **17** | 1900 | 81 | -- | | -- |
| **18** | 1869 | 99 | 65 | biological process assay | -- |
| **19** | 1868 | 106 | 64 | biological process assay | -- |
| **20** | 1867 | 125 | 66 | biological process assay | -- |
| **21** | 1666 | 60 | 58 | protein expression assay | -- |
| **22** | 1666 | 8 | 59 | protein expression assay | -- |
| **23** | 1642 | 26 | 57 | protein expression assay | -- |
| **24** | 1541 | 6 | -- | biological process assay | Other |
| **25** | 1531 | 22 | -- | protein quantification assay | -- |
| **26** | 1524 | 40 | 61 | phosphorylation assay | -- |
| **27** | 1522 | 1450 | 33 | translocation to nucleus assay | -- |
| **28** | 1522 | 62 | 54 | | -- |

| | | | | | |
|---|---|---|---|---|---|
| **29** | 1522 | 30 | 52 | | -- |
| **30** | 1522 | 19 | -- | biological process assay | -- |
| **31** | 1522 | 79 | 54 | | -- |
| **32** | 1522 | 42 | 54 | | -- |
| **33** | 1522 | 14 | 55 | cell viability assay (via ATP quantification) | -- |
| **34** | 1522 | 87 | 60 | protein protein binding assay | Enzyme |
| **35** | 1522 | 14 | 55 | cell viability assay (via ATP quantification) | -- |
| **36** | 1522 | 185 | 56 | | -- |
| **37** | 1522 | 54 | 52 | | -- |
| **38** | 1521 | 149 | -- | biological process assay | -- |
| **39** | 1521 | 29 | -- | biological process assay | -- |
| **40** | 1520 | 88 | -- | biological process assay | -- |
| **41** | 1486 | 513 | 33 | translocation to nucleus assay | -- |
| **42** | 1479 | 40 | -- | | Transcription Regulator |
| **43** | 1396 | 59 | 41 | cell viability assay (via ATP quantification); toxicity assay | -- |
| **44** | 1395 | 55 | 36 | cell viability assay (via ATP quantification); toxicity assay | -- |
| **45** | 1393 | 70 | 35 | cell viability assay (via ATP quantification); toxicity assay | -- |
| **46** | 1392 | 45 | 6 | cell viability assay (via ATP quantification); toxicity assay | -- |
| **47** | 1392 | 57 | 34 | cell viability assay (via ATP quantification); toxicity assay | -- |
| **48** | 1364 | 20 | 25 | gene expression assay | Other |
| **49** | 1364 | 46 | 25 | gene expression assay | Other |
| **50** | 1363 | 18 | 7 | signal transduction assay | -- |
| **51** | 1363 | 26 | 7 | cell viability assay | -- |
| **52** | 1363 | 3 | 40 | signal transduction assay | Other |
| **53** | 1363 | 52 | 32 | protein expression assay | -- |
| **54** | 1363 | 30 | 32 | protein expression assay | -- |
| **55** | 1357 | 88 | 39 | cell viability assay (via ATP quantification) | -- |
| **56** | 1345 | 31 | 32 | signal transduction assay | -- |

| | | | | | |
|---|---|---|---|---|---|
| **57** | 1340 | 94 | 37 | cell viability assay (via ATP quantification) | -- |
| **58** | 1329 | 21 | 33 | translocation to nucleus assay | -- |
| **59** | 1324 | 111 | 50 | signal transduction assay | -- |
| **60** | 1324 | 147 | 50 | signal transduction assay | -- |
| **61** | 1314 | 75 | 38 | cell viability assay (via ATP quantification) | -- |
| **62** | 1310 | 109 | 3 | cell viability assay | Ion Channel |
| **63** | 1310 | 124 | 3 | cell viability assay (via dye reduction) | -- |
| **64** | 1302 | 21 | 7 | protein expression assay | -- |
| **65** | 1302 | 4 | -- | enzyme activity assay | -- |
| **66** | 1302 | 10 | 33 | receptor internalization assay | GPCR |
| **67** | 1302 | 14 | 32 | protein expression assay | -- |
| **68** | 1301 | 14 | 33 | receptor internalization assay | GPCR |
| **69** | 1287 | 122 | 39 | cell viability assay (via ATP quantification) | -- |
| **70** | 1274 | 136 | -- | cell viability assay (via ATP quantification) | -- |
| **71** | 1270 | 36 | -- | | Transcription Regulator |
| **72** | 1268 | 66 | 38 | cell viability assay (via ATP quantification) | -- |
| **73** | 1256 | 74 | 38 | cell viability assay (via ATP quantification) | -- |
| **74** | 1254 | 102 | 37 | cell viability assay (via ATP quantification) | -- |
| **75** | 1236 | 93 | 37 | cell viability assay (via ATP quantification) | -- |
| **76** | 1231 | 65 | 7 | signal transduction assay | -- |
| **77** | 1224 | 46 | 37 | cell viability assay (via ATP quantification) | -- |
| **78** | 1188 | 19 | -- | biological process assay | Transcription Regulator |
| **79** | 1169 | 76 | 39 | cell viability assay (via ATP quantification) | -- |
| **80** | 1164 | 545 | 33 | | -- |
| **81** | 998 | 3 | -- | | -- |
| **82** | 998 | 19 | 43 | cell viability assay (via ATP quantification) | -- |
| **83** | 998 | 34 | 11 | cell viability assay (via ATP quantification) | -- |

| | | | | | |
|---|---|---|---|---|---|
| **84** | 998 | 5 | -- | | -- |
| **85** | 998 | 13 | 9 | cell viability assay (via ATP quantification) | -- |
| **86** | 998 | 75 | 45 | cell viability assay (via ATP quantification) | -- |
| **87** | 998 | 26 | 46 | cell viability assay (via ATP quantification) | -- |
| **88** | 998 | 21 | 46 | cell viability assay (via ATP quantification) | -- |
| **89** | 998 | 18 | 47 | cell viability assay (via ATP quantification) | -- |
| **90** | 998 | 23 | 46 | cell viability assay (via ATP quantification) | -- |
| **91** | 998 | 3 | -- | | -- |
| **92** | 998 | 11 | -- | | -- |
| **93** | 998 | 20 | 11 | | -- |
| **94** | 998 | 14 | 44 | cell viability assay (via ATP quantification) | -- |
| **95** | 998 | 14 | 44 | cell viability assay (via ATP quantification) | -- |
| **96** | 998 | 18 | 47 | cell viability assay (via ATP quantification) | -- |
| **97** | 998 | 985 | 11 | cell viability assay (via ATP quantification) | -- |
| **98** | 998 | 28 | 47 | cell viability assay (via ATP quantification) | -- |
| **99** | 998 | 25 | 9 | cell viability assay (via ATP quantification) | -- |
| **100** | 998 | 24 | 9 | cell viability assay (via ATP quantification) | -- |
| **101** | 998 | 17 | 44 | | -- |
| **102** | 977 | 54 | 38 | cell viability assay (via ATP quantification) | -- |
| **103** | 975 | 23 | 44 | cell viability assay (via ATP quantification) | -- |
| **104** | 975 | 66 | 44 | cell viability assay (via ATP quantification) | -- |
| **105** | 975 | 936 | 44 | cell viability assay (via ATP quantification) | -- |
| **106** | 975 | 34 | 44 | cell viability assay (via ATP quantification) | -- |
| **107** | 975 | 50 | 44 | cell viability assay (via ATP quantification) | -- |
| **108** | 971 | 176 | 44 | cell viability assay (via ATP quantification) | -- |
| **109** | 932 | 19 | 49 | protein stability assay | -- |

| | | | | | |
|---|---|---|---|---|---|
| **110** | 932 | 31 | 48 | cell viability assay (via ATP quantification) | -- |
| **111** | 916 | 2 | -- | enzyme activity assay | Enzyme |
| **112** | 851 | 851 | -- | cell differentiation assay | -- |
| **113** | 835 | 104 | 39 | cell viability assay (via ATP quantification) | -- |
| **114** | 791 | 37 | 3 | signal transduction assay | Enzyme |
| **115** | 717 | 2 | 3 | protein binding assay | -- |
| **116** | 717 | 1 | 25 | protein binding assay | -- |
| **117** | 717 | 54 | 1 | toxicity assay | GPCR |
| **118** | 717 | 54 | 0 | toxicity assay | GPCR |
| **119** | 717 | 27 | -- | | -- |
| **120** | 717 | 11 | 3 | protein binding assay | -- |
| **121** | 717 | 16 | 3 | protein binding assay | -- |
| **122** | 717 | 36 | 25 | protein binding assay | -- |
| **123** | 717 | 52 | 1 | toxicity assay | GPCR |
| **124** | 717 | 7 | 25 | protein binding assay | Kinase |
| **125** | 717 | 6 | 3 | protein binding assay | -- |
| **126** | 716 | 52 | 0 | toxicity assay | GPCR |
| **127** | 715 | 37 | 25 | protein binding assay | -- |
| **128** | 715 | 7 | 25 | protein binding assay | -- |
| **129** | 715 | 43 | 25 | protein binding assay | -- |
| **130** | 715 | 12 | 25 | protein binding assay | -- |
| **131** | 715 | 1 | 67 | protein binding assay | -- |
| **132** | 715 | 1 | 67 | protein binding assay | -- |
| **133** | 715 | 3 | 67 | protein binding assay | -- |
| **134** | 715 | 3 | 3 | protein binding assay | -- |
| **135** | 715 | 1 | 67 | protein binding assay | |
| **136** | 715 | 3 | 3 | protein binding assay | -- |
| **137** | 711 | 8 | 3 | protein binding assay | -- |
| **138** | 708 | 6 | 25 | protein binding assay | -- |
| **139** | 706 | 69 | 3 | protein binding assay | Transcription Regulator |
| **140** | 705 | 118 | 3 | protein binding assay | -- |
| **141** | 642 | 5 | 3 | protein binding assay | -- |
| **142** | 642 | 7 | 3 | protein binding assay | -- |
| **143** | 642 | 5 | 3 | protein binding assay | -- |
| **144** | 642 | 3 | 3 | protein binding assay | -- |
| **145** | 641 | 8 | 3 | protein binding assay | -- |
| **146** | 641 | 12 | 3 | protein binding assay | -- |
| **147** | 641 | 3 | 3 | protein binding assay | -- |

| | | | | | |
|---|---|---|---|---|---|
| **148** | 638 | 26 | 3 | protein binding assay | -- |
| **149** | 625 | 80 | 3 | protein binding assay | -- |
| **150** | 558 | 63 | 20 | cell proliferation assay | -- |
| **151** | 558 | 43 | -- | toxicity assay | -- |
| **152** | 554 | 62 | 20 | cell proliferation assay | -- |
| **153** | 554 | 36 | 20 | cell proliferation assay | -- |
| **154** | 540 | 49 | 20 | cell proliferation assay | -- |
| **155** | 506 | 47 | 14 | cell proliferation assay | -- |
| **156** | 506 | 12 | 51 | cell proliferation assay | -- |
| **157** | 506 | 12 | 51 | cell proliferation assay | Enzyme |
| **158** | 506 | 20 | 51 | cell proliferation assay | -- |
| **159** | 506 | 20 | 53 | cell proliferation assay | -- |
| **160** | 506 | 23 | 51 | | -- |
| **161** | 499 | 8 | -- | cell proliferation assay | -- |
| **162** | 498 | 29 | 5 | cell proliferation assay | -- |
| **163** | 497 | 45 | -- | | -- |
| **164** | 497 | 106 | -- | | -- |
| **165** | 497 | 3 | -- | cell proliferation assay | -- |
| **166** | 497 | 15 | -- | cell proliferation assay | -- |
| **167** | 497 | 4 | -- | cell proliferation assay | -- |
| **168** | 497 | 3 | -- | cell proliferation assay | -- |
| **169** | 497 | 2 | -- | cell proliferation assay | -- |
| **170** | 497 | 7 | -- | cell proliferation assay | -- |
| **171** | 497 | 4 | -- | cell proliferation assay | -- |
| **172** | 497 | 72 | -- | | -- |
| **173** | 496 | 73 | -- | | -- |
| **174** | 495 | 29 | -- | cell proliferation assay | -- |
| **175** | 491 | 26 | 9 | cell proliferation assay | -- |
| **176** | 486 | 38 | -- | | -- |
| **177** | 486 | 36 | 3 | protein binding assay | -- |
| **178** | 486 | 54 | 3 | protein binding assay | -- |
| **179** | 484 | 40 | -- | protein binding assay | -- |
| **180** | 478 | 22 | -- | | -- |
| **181** | 478 | 15 | -- | | -- |
| **182** | 477 | 26 | 42 | cell proliferation assay | Kinase |
| **183** | 477 | 58 | -- | | -- |
| **184** | 477 | 29 | 42 | cell proliferation assay | Kinase |
| **185** | 476 | 22 | 42 | cell proliferation assay | Kinase |
| **186** | 476 | 26 | 42 | cell proliferation assay | Kinase |
| **187** | 464 | 36 | 42 | cell proliferation assay | Kinase |

| | | | | | |
|---|---|---|---|---|---|
| **188** | 436 | 30 | 17 | cell proliferation assay | -- |
| **189** | 436 | 20 | -- | cell proliferation assay | -- |
| **190** | 436 | 17 | 10 | cell proliferation assay | -- |
| **191** | 436 | 39 | 7 | cell proliferation assay | -- |
| **192** | 434 | 22 | 18 | cell proliferation assay | -- |
| **193** | 434 | 28 | -- | cell proliferation assay | Enzyme |
| **194** | 434 | 28 | 15 | cell proliferation assay | -- |
| **195** | 434 | 35 | 19 | cell proliferation assay | -- |
| **196** | 434 | 24 | -- | cell proliferation assay | -- |
| **197** | 433 | 29 | 21 | | -- |
| **198** | 433 | 31 | 20 | toxicity assay | -- |
| **199** | 433 | 26 | 8 | cell proliferation assay | Kinase |
| **200** | 433 | 28 | -- | cell proliferation assay | -- |
| **201** | 429 | 13 | 31 | cell proliferation assay | -- |
| **202** | 429 | 20 | 30 | cell proliferation assay | -- |
| **203** | 429 | 21 | 30 | cell proliferation assay | -- |
| **204** | 429 | 10 | -- | | -- |
| **205** | 429 | 5 | 11 | cell proliferation assay | -- |
| **206** | 429 | 203 | 9 | gene expression assay | -- |
| **207** | 429 | 38 | 28 | cell proliferation assay | Kinase |
| **208** | 429 | 13 | -- | | -- |
| **209** | 429 | 24 | 29 | cell proliferation assay | -- |
| **210** | 429 | 25 | 29 | cell proliferation assay | -- |
| **211** | 429 | 8 | -- | cell proliferation assay | -- |
| **212** | 429 | 21 | -- | cell proliferation assay | -- |
| **213** | 429 | 45 | -- | | -- |
| **214** | 429 | 12 | 26 | | -- |
| **215** | 429 | 19 | -- | | -- |
| **216** | 429 | 28 | 28 | cell proliferation assay | Kinase |
| **217** | 429 | 18 | 31 | cell proliferation assay | -- |
| **218** | 429 | 30 | 28 | cell proliferation assay | Kinase |
| **219** | 429 | 14 | -- | | -- |
| **220** | 429 | 18 | -- | cell proliferation assay | -- |
| **221** | 429 | 17 | -- | | -- |
| **222** | 429 | 17 | -- | cell proliferation assay | -- |
| **223** | 428 | 22 | 23 | | -- |
| **224** | 428 | 29 | 25 | calcium quantification assay | -- |
| **225** | 428 | 26 | 28 | cell proliferation assay | Kinase |
| **226** | 428 | 56 | 27 | toxicity assay | -- |
| **227** | 428 | 49 | -- | toxicity assay | -- |

| | | | | | |
|---|---|---|---|---|---|
| **228** | 426 | 15 | 6 | cell proliferation assay | -- |
| **229** | 426 | 65 | -- | cell proliferation assay | -- |
| **230** | 426 | 22 | -- | cell proliferation assay | -- |
| **231** | 426 | 23 | -- | cell proliferation assay | -- |
| **232** | 426 | 24 | 14 | cell proliferation assay | -- |
| **233** | 426 | 17 | -- | cell proliferation assay | -- |
| **234** | 426 | 24 | 5 | cell proliferation assay | -- |
| **235** | 425 | 13 | -- | | -- |
| **236** | 425 | 36 | 16 | cell proliferation assay | -- |
| **237** | 425 | 108 | -- | toxicity assay | -- |
| **238** | 425 | 20 | 22 | | -- |
| **239** | 425 | 34 | -- | | -- |
| **240** | 425 | 44 | -- | cell proliferation assay | -- |
| **241** | 425 | 15 | 22 | | -- |
| **242** | 425 | 24 | -- | cell proliferation assay | -- |
| **243** | 424 | 20 | 12 | | -- |
| **244** | 424 | 21 | 11 | | -- |
| **245** | 424 | 65 | 12 | | -- |
| **246** | 424 | 12 | 13 | cell proliferation assay | -- |
| **247** | 424 | 23 | 13 | cell proliferation assay | -- |
| **248** | 424 | 27 | 11 | | -- |
| **249** | 424 | 43 | 11 | | -- |
| **250** | 424 | 15 | 13 | cell proliferation assay | -- |
| **251** | 423 | 32 | 12 | | -- |
| **252** | 423 | 34 | 13 | cell proliferation assay | -- |
| **253** | 420 | 42 | -- | cell proliferation assay | -- |
| **254** | 418 | 40 | -- | | -- |
| **255** | 415 | 45 | 4 | cell proliferation assay | -- |
| **256** | 413 | 77 | -- | cell differentiation assay | -- |
| **257** | 410 | 27 | -- | cell proliferation assay | -- |
| **258** | 410 | 63 | 24 | | -- |
| **259** | 409 | 6 | -- | | -- |
| **260** | 408 | 39 | -- | cell proliferation assay | -- |
| **261** | 403 | 71 | 9 | toxicity assay | Other |

| Assay ID | No. of compounds | No. of active compounds | Cell line | Assay type | Target class |
|---|---|---|---|---|---|
| **0** | 2248 | 36 | | biological process assay | Protease |

| | | | | | |
|---|---|---|---|---|---|
| **1** | 2248 | 149 | | | -- |
| **2** | 2248 | 153 | | | -- |
| **3** | 2248 | 191 | | | -- |
| **4** | 2248 | 1420 | | cell viability assay (via ATP quantification) | Kinase |
| **5** | 2248 | 253 | | | -- |
| **6** | 2248 | 89 | | biological process assay | Other |
| **7** | 2248 | 38 | | biological process assay | -- |
| **8** | 2247 | 120 | 0 | biological process assay | -- |
| **9** | 2245 | 11 | 1 | cell viability assay (via ATP quantification); necrosis assay | -- |
| **10** | 2017 | 7 | 2 | cAMP quantification assay | -- |
| **11** | 2017 | 18 | 3 | enzyme activity assay | Enzyme |
| **12** | 1928 | 133 | 4 | | -- |
| **13** | 1928 | 110 | 4 | cell viability assay | -- |
| **14** | 1900 | 65 | | | -- |
| **15** | 1900 | 187 | 5 | protein quantification assay | -- |
| **16** | 1900 | 8 | | protein degradation assay; protein protein binding assay | -- |
| **17** | 1900 | 81 | | | -- |
| **18** | 1869 | 99 | 6 | biological process assay | -- |
| **19** | 1868 | 106 | 7 | biological process assay | -- |
| **20** | 1867 | 125 | 8 | biological process assay | -- |
| **21** | 1666 | 60 | 9 | protein expression assay | -- |
| **22** | 1666 | 8 | 4 | protein expression assay | -- |
| **23** | 1642 | 26 | 10 | protein expression assay | -- |
| **24** | 1541 | 6 | | biological process assay | Other |
| **25** | 1531 | 22 | | protein quantification assay | -- |
| **26** | 1524 | 40 | 11 | phosphorylation assay | -- |
| **27** | 1522 | 1450 | 12 | translocation to nucleus assay | -- |
| **28** | 1522 | 62 | 13 | | -- |
| **29** | 1522 | 30 | 14 | | -- |
| **30** | 1522 | 19 | | biological process assay | -- |
| **31** | 1522 | 79 | 13 | | -- |
| **32** | 1522 | 42 | 13 | | -- |
| **33** | 1522 | 14 | 15 | cell viability assay (via ATP quantification) | -- |

| | | | | | |
|---|---|---|---|---|---|
| **34** | 1522 | 87 | 16 | protein protein binding assay | Enzyme |
| **35** | 1522 | 14 | 15 | cell viability assay (via ATP quantification) | -- |
| **36** | 1522 | 185 | 17 | | -- |
| **37** | 1522 | 54 | 14 | | -- |
| **38** | 1521 | 149 | | biological process assay | -- |
| **39** | 1521 | 29 | | biological process assay | -- |
| **40** | 1520 | 88 | | biological process assay | -- |
| **41** | 1486 | 513 | 12 | translocation to nucleus assay | -- |
| **42** | 1479 | 40 | | | Transcription Regulator |
| **43** | 1396 | 59 | 18 | cell viability assay (via ATP quantification); toxicity assay | -- |
| **44** | 1395 | 55 | 19 | cell viability assay (via ATP quantification); toxicity assay | -- |
| **45** | 1393 | 70 | 20 | cell viability assay (via ATP quantification); toxicity assay | -- |
| **46** | 1392 | 45 | 21 | cell viability assay (via ATP quantification); toxicity assay | -- |
| **47** | 1392 | 57 | 22 | cell viability assay (via ATP quantification); toxicity assay | -- |
| **48** | 1364 | 20 | 2 | gene expression assay | Other |
| **49** | 1364 | 46 | 2 | gene expression assay | Other |
| **50** | 1363 | 18 | 23 | signal transduction assay | -- |
| **51** | 1363 | 26 | 23 | cell viability assay | -- |
| **52** | 1363 | 3 | 24 | signal transduction assay | Other |
| **53** | 1363 | 52 | 25 | protein expression assay | -- |
| **54** | 1363 | 30 | 25 | protein expression assay | -- |
| **55** | 1357 | 88 | 26 | cell viability assay (via ATP quantification) | -- |
| **56** | 1345 | 31 | 25 | signal transduction assay | -- |
| **57** | 1340 | 94 | 27 | cell viability assay (via ATP quantification) | -- |
| **58** | 1329 | 21 | 12 | translocation to nucleus assay | -- |
| **59** | 1324 | 111 | 28 | signal transduction assay | -- |
| **60** | 1324 | 147 | 28 | signal transduction assay | -- |

| | | | | | |
|---|---|---|---|---|---|
| **61** | 1314 | 75 | 29 | cell viability assay (via ATP quantification) | -- |
| **62** | 1310 | 109 | 30 | cell viability assay | Ion Channel |
| **63** | 1310 | 124 | 30 | cell viability assay (via dye reduction) | -- |
| **64** | 1302 | 21 | 23 | protein expression assay | -- |
| **65** | 1302 | 4 | | enzyme activity assay | -- |
| **66** | 1302 | 10 | 12 | receptor internalization assay | GPCR |
| **67** | 1302 | 14 | 25 | protein expression assay | -- |
| **68** | 1301 | 14 | 12 | receptor internalization assay | GPCR |
| **69** | 1287 | 122 | 26 | cell viability assay (via ATP quantification) | -- |
| **70** | 1274 | 136 | | cell viability assay (via ATP quantification) | -- |
| **71** | 1270 | 36 | | | Transcription Regulator |
| **72** | 1268 | 66 | 29 | cell viability assay (via ATP quantification) | -- |
| **73** | 1256 | 74 | 29 | cell viability assay (via ATP quantification) | -- |
| **74** | 1254 | 102 | 27 | cell viability assay (via ATP quantification) | -- |
| **75** | 1236 | 93 | 27 | cell viability assay (via ATP quantification) | -- |
| **76** | 1231 | 65 | 23 | signal transduction assay | -- |
| **77** | 1224 | 46 | 27 | cell viability assay (via ATP quantification) | -- |
| **78** | 1188 | 19 | | biological process assay | Transcription Regulator |
| **79** | 1169 | 76 | 26 | cell viability assay (via ATP quantification) | -- |
| **80** | 1164 | 545 | 12 | | -- |
| **81** | 998 | 3 | | | -- |
| **82** | 998 | 19 | 31 | cell viability assay (via ATP quantification) | -- |
| **83** | 998 | 34 | 32 | cell viability assay (via ATP quantification) | -- |
| **84** | 998 | 5 | | | -- |
| **85** | 998 | 13 | 33 | cell viability assay (via ATP quantification) | -- |
| **86** | 998 | 75 | 34 | cell viability assay (via ATP quantification) | -- |

| | | | | | |
|---|---|---|---|---|---|
| **87** | 998 | 26 | 35 | cell viability assay (via ATP quantification) | -- |
| **88** | 998 | 21 | 35 | cell viability assay (via ATP quantification) | -- |
| **89** | 998 | 18 | 36 | cell viability assay (via ATP quantification) | -- |
| **90** | 998 | 23 | 35 | cell viability assay (via ATP quantification) | -- |
| **91** | 998 | 3 | | | -- |
| **92** | 998 | 11 | | | -- |
| **93** | 998 | 20 | 32 | | -- |
| **94** | 998 | 14 | 37 | cell viability assay (via ATP quantification) | -- |
| **95** | 998 | 14 | 37 | cell viability assay (via ATP quantification) | -- |
| **96** | 998 | 18 | 36 | cell viability assay (via ATP quantification) | -- |
| **97** | 998 | 985 | 32 | cell viability assay (via ATP quantification) | -- |
| **98** | 998 | 28 | 36 | cell viability assay (via ATP quantification) | -- |
| **99** | 998 | 25 | 33 | cell viability assay (via ATP quantification) | -- |
| **100** | 998 | 24 | 33 | cell viability assay (via ATP quantification) | -- |
| **101** | 998 | 17 | 37 | | -- |
| **102** | 977 | 54 | 29 | cell viability assay (via ATP quantification) | -- |
| **103** | 975 | 23 | 37 | cell viability assay (via ATP quantification) | -- |
| **104** | 975 | 66 | 37 | cell viability assay (via ATP quantification) | -- |
| **105** | 975 | 936 | 37 | cell viability assay (via ATP quantification) | -- |
| **106** | 975 | 34 | 37 | cell viability assay (via ATP quantification) | -- |
| **107** | 975 | 50 | 37 | cell viability assay (via ATP quantification) | -- |
| **108** | 971 | 176 | 37 | cell viability assay (via ATP quantification) | -- |
| **109** | 932 | 19 | 38 | protein stability assay | -- |
| **110** | 932 | 31 | 39 | cell viability assay (via ATP quantification) | -- |
| **111** | 916 | 2 | | enzyme activity assay | Enzyme |
| **112** | 851 | 851 | | cell differentiation assay | -- |

| | | | | | |
|---|---|---|---|---|---|
| **113** | 835 | 104 | 26 | cell viability assay (via ATP quantification) | -- |
| **114** | 791 | 37 | 30 | signal transduction assay | Enzyme |
| **115** | 717 | 2 | 30 | protein binding assay | -- |
| **116** | 717 | 1 | 2 | protein binding assay | -- |
| **117** | 717 | 54 | 40 | toxicity assay | GPCR |
| **118** | 717 | 54 | 41 | toxicity assay | GPCR |
| **119** | 717 | 27 | | | -- |
| **120** | 717 | 11 | 30 | protein binding assay | -- |
| **121** | 717 | 16 | 30 | protein binding assay | -- |
| **122** | 717 | 36 | 2 | protein binding assay | -- |
| **123** | 717 | 52 | 40 | toxicity assay | GPCR |
| **124** | 717 | 7 | 2 | protein binding assay | Kinase |
| **125** | 717 | 6 | 30 | protein binding assay | -- |
| **126** | 716 | 52 | 41 | toxicity assay | GPCR |
| **127** | 715 | 37 | 2 | protein binding assay | -- |
| **128** | 715 | 7 | 2 | protein binding assay | -- |
| **129** | 715 | 43 | 2 | protein binding assay | -- |
| **130** | 715 | 12 | 2 | protein binding assay | -- |
| **131** | 715 | 1 | 42 | protein binding assay | -- |
| **132** | 715 | 1 | 42 | protein binding assay | -- |
| **133** | 715 | 3 | 42 | protein binding assay | -- |
| **134** | 715 | 3 | 30 | protein binding assay | -- |
| **135** | 715 | 1 | 42 | protein binding assay | |
| **136** | 715 | 3 | 30 | protein binding assay | -- |
| **137** | 711 | 8 | 30 | protein binding assay | -- |
| **138** | 708 | 6 | 2 | protein binding assay | -- |
| **139** | 706 | 69 | 30 | protein binding assay | Transcription Regulator |
| **140** | 705 | 118 | 30 | protein binding assay | -- |
| **141** | 642 | 5 | 30 | protein binding assay | -- |
| **142** | 642 | 7 | 30 | protein binding assay | -- |
| **143** | 642 | 5 | 30 | protein binding assay | -- |
| **144** | 642 | 3 | 30 | protein binding assay | -- |
| **145** | 641 | 8 | 30 | protein binding assay | -- |
| **146** | 641 | 12 | 30 | protein binding assay | -- |
| **147** | 641 | 3 | 30 | protein binding assay | -- |
| **148** | 638 | 26 | 30 | protein binding assay | -- |
| **149** | 625 | 80 | 30 | protein binding assay | -- |
| **150** | 558 | 63 | 43 | cell proliferation assay | -- |

| | | | | | |
|---|---|---|---|---|---|
| **151** | 558 | 43 | | toxicity assay | -- |
| **152** | 554 | 62 | 43 | cell proliferation assay | -- |
| **153** | 554 | 36 | 43 | cell proliferation assay | -- |
| **154** | 540 | 49 | 43 | cell proliferation assay | -- |
| **155** | 506 | 47 | 44 | cell proliferation assay | -- |
| **156** | 506 | 12 | 45 | cell proliferation assay | -- |
| **157** | 506 | 12 | 45 | cell proliferation assay | Enzyme |
| **158** | 506 | 20 | 45 | cell proliferation assay | -- |
| **159** | 506 | 20 | 46 | cell proliferation assay | -- |
| **160** | 506 | 23 | 45 | | -- |
| **161** | 499 | 8 | | cell proliferation assay | -- |
| **162** | 498 | 29 | 47 | cell proliferation assay | -- |
| **163** | 497 | 45 | | | -- |
| **164** | 497 | 106 | | | -- |
| **165** | 497 | 3 | | cell proliferation assay | -- |
| **166** | 497 | 15 | | cell proliferation assay | -- |
| **167** | 497 | 4 | | cell proliferation assay | -- |
| **168** | 497 | 3 | | cell proliferation assay | -- |
| **169** | 497 | 2 | | cell proliferation assay | -- |
| **170** | 497 | 7 | | cell proliferation assay | -- |
| **171** | 497 | 4 | | cell proliferation assay | -- |
| **172** | 497 | 72 | | | -- |
| **173** | 496 | 73 | | | -- |
| **174** | 495 | 29 | | cell proliferation assay | -- |
| **175** | 491 | 26 | 33 | cell proliferation assay | -- |
| **176** | 486 | 38 | | | -- |
| **177** | 486 | 36 | 30 | protein binding assay | -- |
| **178** | 486 | 54 | 30 | protein binding assay | -- |
| **179** | 484 | 40 | | protein binding assay | -- |
| **180** | 478 | 22 | | | -- |
| **181** | 478 | 15 | | | -- |
| **182** | 477 | 26 | 48 | cell proliferation assay | Kinase |
| **183** | 477 | 58 | | | -- |
| **184** | 477 | 29 | 48 | cell proliferation assay | Kinase |
| **185** | 476 | 22 | 48 | cell proliferation assay | Kinase |
| **186** | 476 | 26 | 48 | cell proliferation assay | Kinase |
| **187** | 464 | 36 | 48 | cell proliferation assay | Kinase |
| **188** | 436 | 30 | 49 | cell proliferation assay | -- |
| **189** | 436 | 20 | | cell proliferation assay | -- |

| | | | | | |
|---|---|---|---|---|---|
| **190** | 436 | 17 | 50 | cell proliferation assay | -- |
| **191** | 436 | 39 | 23 | cell proliferation assay | -- |
| **192** | 434 | 22 | 51 | cell proliferation assay | -- |
| **193** | 434 | 28 | | cell proliferation assay | Enzyme |
| **194** | 434 | 28 | 52 | cell proliferation assay | -- |
| **195** | 434 | 35 | 53 | cell proliferation assay | -- |
| **196** | 434 | 24 | | cell proliferation assay | -- |
| **197** | 433 | 29 | 54 | | -- |
| **198** | 433 | 31 | 43 | toxicity assay | -- |
| **199** | 433 | 26 | 55 | cell proliferation assay | Kinase |
| **200** | 433 | 28 | | cell proliferation assay | -- |
| **201** | 429 | 13 | 56 | cell proliferation assay | -- |
| **202** | 429 | 20 | 57 | cell proliferation assay | -- |
| **203** | 429 | 21 | 57 | cell proliferation assay | -- |
| **204** | 429 | 10 | | | -- |
| **205** | 429 | 5 | 32 | cell proliferation assay | -- |
| **206** | 429 | 203 | 33 | gene expression assay | -- |
| **207** | 429 | 38 | 58 | cell proliferation assay | Kinase |
| **208** | 429 | 13 | | | -- |
| **209** | 429 | 24 | 59 | cell proliferation assay | -- |
| **210** | 429 | 25 | 59 | cell proliferation assay | -- |
| **211** | 429 | 8 | | cell proliferation assay | -- |
| **212** | 429 | 21 | | cell proliferation assay | -- |
| **213** | 429 | 45 | | | -- |
| **214** | 429 | 12 | 60 | | -- |
| **215** | 429 | 19 | | | -- |
| **216** | 429 | 28 | 58 | cell proliferation assay | Kinase |
| **217** | 429 | 18 | 56 | cell proliferation assay | -- |
| **218** | 429 | 30 | 58 | cell proliferation assay | Kinase |
| **219** | 429 | 14 | | | -- |
| **220** | 429 | 18 | | cell proliferation assay | -- |
| **221** | 429 | 17 | | | -- |
| **222** | 429 | 17 | | cell proliferation assay | -- |
| **223** | 428 | 22 | 61 | | -- |
| **224** | 428 | 29 | 2 | calcium quantification assay | -- |
| **225** | 428 | 26 | 58 | cell proliferation assay | Kinase |
| **226** | 428 | 56 | 62 | toxicity assay | -- |
| **227** | 428 | 49 | | toxicity assay | -- |
| **228** | 426 | 15 | 21 | cell proliferation assay | -- |

| | | | | | |
|---|---|---|---|---|---|
| **229** | 426 | 65 | | cell proliferation assay | -- |
| **230** | 426 | 22 | | cell proliferation assay | -- |
| **231** | 426 | 23 | | cell proliferation assay | -- |
| **232** | 426 | 24 | 44 | cell proliferation assay | -- |
| **233** | 426 | 17 | | cell proliferation assay | -- |
| **234** | 426 | 24 | 47 | cell proliferation assay | -- |
| **235** | 425 | 13 | | | -- |
| **236** | 425 | 36 | 63 | cell proliferation assay | -- |
| **237** | 425 | 108 | | toxicity assay | -- |
| **238** | 425 | 20 | 64 | | -- |
| **239** | 425 | 34 | | | -- |
| **240** | 425 | 44 | | cell proliferation assay | -- |
| **241** | 425 | 15 | 64 | | -- |
| **242** | 425 | 24 | | cell proliferation assay | -- |
| **243** | 424 | 20 | 65 | | -- |
| **244** | 424 | 21 | 32 | | -- |
| **245** | 424 | 65 | 65 | | -- |
| **246** | 424 | 12 | 66 | cell proliferation assay | -- |
| **247** | 424 | 23 | 66 | cell proliferation assay | -- |
| **248** | 424 | 27 | 32 | | -- |
| **249** | 424 | 43 | 32 | | -- |
| **250** | 424 | 15 | 66 | cell proliferation assay | -- |
| **251** | 423 | 32 | 65 | | -- |
| **252** | 423 | 34 | 66 | cell proliferation assay | -- |
| **253** | 420 | 42 | | cell proliferation assay | -- |
| **254** | 418 | 40 | | | -- |
| **255** | 415 | 45 | 67 | cell proliferation assay | -- |
| **256** | 413 | 77 | | cell differentiation assay | -- |
| **257** | 410 | 27 | | cell proliferation assay | -- |
| **258** | 410 | 63 | 68 | | -- |
| **259** | 409 | 6 | | | -- |
| **260** | 408 | 39 | | cell proliferation assay | -- |
| **261** | 403 | 71 | 33 | toxicity assay | Other |
